# Combined production of Non-Hemolytic Enterotoxin and Sphingomyelinase as a marker of diarrheal food poisoning strains in the *Bacillus cereus* group

**DOI:** 10.64898/2026.08.27.747690

**Authors:** Priscilla F. Cardoso, Nathalie Gilois, Gislayne T. Vilas-Bôas, Didier Lereclus, Michel Gohar, Stéphane Perchat, Leyla Slamti

## Abstract

The *Bacillus cereus* group comprises bacteria of biotechnological interest, but also raises health concerns. Some bacteria in this group are opportunistic human pathogens, mainly causing foodborne gastrointestinal infections. As of today, the presence, sequence variability, or expression of genes encoding toxins or other virulence factors are insufficient to predict the potential of a given isolate to cause the diarrheal form of the disease. To address this limitation, we developed a sandwich ELISA to quantify the NheA and Sphingomyelinase (SMase) proteins in culture supernatants to test them as markers of pathogenic potential. Application of the assay to a collection of *B. cereus* group isolates revealed that strains associated with food poisoning outbreaks produce significantly more NheA and SMase than those isolated from the environment or from commercial products. Statistical analyses show that the combined quantification of NheA and SMase provides robust discrimination between pathogenic and non-pathogenic (environmental and commercial) profiles. These results demonstrate that the quantitative assessment of both NheA and SMase production can serve as a reliable biomarker for distinguishing diarrheic food poisoning isolates from harmless strains.

## Introduction

The *Bacillus cereus* group comprises bacteria of clinical and economic relevance that are closely related genetically, although presenting different pathogenic profiles. Among the main species of the group are *Bacillus thuringiensis*, an insect pathogen of commercial and biotechnological interest used worldwide as a biopesticide; *Bacillus anthracis*, the etiological agent of anthrax; and *Bacillus cereus*, an opportunistic pathogen often involved in food poisoning outbreaks [1,2]. There are two different types of foodborne diseases caused by *B. cereus* group strains: an emetic form, caused by the cereulide toxin, and a diarrheic form, whose mode of action is not completely understood [3,4]. Taking this into account, many efforts have been made to develop methods for distinguishing innocuous *B. cereus* strains from pathogenic ones, such as phylogenetic analysis or the identification of pathogenic markers.

Genomic assessment of *B. cereus* group bacteria leads to the classification of the isolates into seven phylogenetic groups, determined by the genetic sequences of housekeeping genes (such as *panC*) [5]. This classification generally assembles together phenotypically similar strains, but not exclusively. Indeed, Group II is composed of psychrotolerant strains, including some associated with food poisoning, *B. anthracis* and other human pathogenic strains are found in Group III, *B. thuringiensis* and environmental isolates in Group IV, *B. toyonensis* in Group V, and psychrotrophic strains like *Bacillus mycoides* and *Bacillus weihenstephanensis* in Group VI. It was shown that this classification correlates relatively well with the ecological diversification or pathogenic potential of the group [5,6].

Specific virulence factor determinants, such as genes encoding *B. thuringiensis* insecticidal proteins, *B. anthracis* capsule and toxins, and the non-ribosomal peptide synthase responsible for the production of cereulide (the emetic toxin), are found on plasmids [2]. These plasmids can be transferred among bacteria, but the presence of these genes can serve as a molecular marker to detect isolates that could produce these toxins. In sharp contrast, there is still no specific pathogenic trait clearly associated with diarrhea-causing strains [2,7,8]. Consequently, it is difficult to predict the diarrheal potential of a new isolate belonging to the *B. cereus* group, and the identification of bacteria able to cause the diarrheic form of the foodborne illness is far more complex.

Based on their biological properties (protease, phospholipase, cytotoxin, etc.), several proteins can be regarded as potential virulence factors that may contribute to diarrheal syndromes. Among such factors, several have been previously investigated; including the two tripartite enterotoxins hemolysin BL (Hbl) and nonhemolytic enterotoxin (Nhe), and the single-protein cytotoxin K (CytK) [2,4]. However, the presence or absence of their encoding genes did not correlate with the pathogenic profile of the isolates, since Hbl and CytK are absent in strains related to severe cases of food poisoning outbreaks, such as strain NVH0075/95 [9]. In contrast, the Nhe-encoding genes are found in virtually all *B. cereus* group strains. Other virulence factors include a wide range of secreted degradative enzymes, such as the phospholipases phosphatidylinositol-specific phospholipase C (PI-PLC), phosphatidylcholine-specific phospholipase C (PC-PLC), and sphingomyelinase (SMase).

All the genes encoding these potential virulence factors have a chromosomal location and their transcription is positively activated by the virulence regulator PlcR [10]. This regulation enables bacteria to adjust gene expression to population density, since PlcR is part of a quorum-sensing system and is activated by its cognate peptide PapR [11]. The PlcR-PapR system controls the expression of at least 45 genes, most of which encode extracellular proteins, such as the potential virulence factors presented above [12,13]. Overall, the key role of the PlcR regulon in bacterial virulence has been demonstrated *in vivo* in insects, mice, and rabbits by assaying the pathogenicity of *plcR*-deficient strains [14,15]. However, the transcription of genes encoding these virulence factors alone provides limited information regarding the pathogenic potential of a *B. cereus* isolate. Indeed, only partial correlation has been observed between the expression levels and the production titers of virulence factors [9,16]. Thus, the production of virulence factors might be subject to other regulatory mechanisms at post-transcriptional, post-translational, and secretion levels [9,17–19]. Hence, it has been suggested that the pathogenicity linked to the diarrheic disease could be multifactorial and related to the quantity of toxins and other virulence factors produced [9,20]. However, no animal model is available to confirm the involvement of each virulence factor in diarrheal syndromes.

The interaction of different virulence factors could be required to establish infection and pathogenicity. For example, Nhe and SMase were shown to cooperate in cytotoxicity and pathogenicity of *B. cereus*, highlighting their role in the virulence of these bacteria [21]. The Nhe toxin is composed of NheA (41.0 kDa), NheB (39.8 kDa), and NheC (36.5 kDa), which are encoded by the operon including the *nheA*, *nheB*, and *nheC* genes, respectively [22,23]. This toxin was first described in the strain NVH0075/95, isolated from a foodborne outbreak in Norway [24]. The three Nhe components exhibit homology among themselves and also with Hbl components, both belonging to the ClyA family of pore-forming toxins. All three Nhe proteins are needed for the full activity of the toxin [23–25]. SMase (34 kDa) was shown to play a role in the cytotoxicity and pathogenicity of *B. cereus* group bacteria, helping bacteria to evade the host innate immune response [21,26]. SMase hydrolyses sphingomyelin in phosphocholine and ceramide and is encoded by the *sph* gene found in an operon, downstream from the *plcB* gene, which encodes PC-PLC. These two enzymes (SMase and PC-PLC) work together to elicit cytolytic activity and were first described as the complex cereolysin AB [27]. In addition, SMase also acts synergistically with Hbl to enhance its hemolytic activity [28]. Taking into account the interaction of these virulence factors and the fact that the production of SMase was shown to be greater in clinical isolates than in environmental ones [26], SMase appears to play an important role in *B. cereus* pathogenesis. This role in pathogenesis has already been described for structurally related proteins from other bacteria, such as *Staphylococcus aureus*, *Clostridium perfringens*, and *Listeria ivanovii* [29].

Considering the absence of a molecular marker that enables easy distinction between diarrheic food poisoning *B. cereus* isolates from innocuous strains, we developed an immunodetection quantitative assay to precisely quantify the production of NheA and SMase by strains of the *B. cereus* group. We found that it is possible to distinguish foodborne pathogenic strains from nonpathogenic ones (environmental and commercial) using the method described in this study.

## Materials and Methods

### 1. Bacterial strains and culture conditions

Table 1 presents the list of the *B. cereus* group strains used in this study. We selected 25 strains divided into three profiles: 10 strains classified as Environmental, gathering strains associated to processed or non-processed food and laboratory environment; five strains categorized as Commercial, isolated from biopesticide and probiotic products; and 10 defined as Food Poisoning, isolated from diarrheal outbreaks. The strains NVH0075-95, ATCC14579, Bt407, and Bt407Δ*plcR* were used as controls for test development. To evaluate the production of NheA and SMase, the strains were cultivated in LB medium at 37°C until two hours after the entry into the stationary phase, a growth stage where both NheA and SMase were found in high concentration in the culture supernatant [13,30]. Collected samples were then centrifuged at 13000 rpm for 3 min.; cell-free supernatants were obtained by filtration using 0.22µm PVDF syringe filters (Merck Millipore) and stored at-20°C until use. *E. coli* K-12 strain TG1 and *E. coli* M15 [pREP4] recombinant strains were cultivated in LB medium at 30 or 37°C, and, when required, ampicillin 100 μg/mL and kanamycin 25 μg/mL were added.

**Table 1.** Strains of the *B. cereus* group used in this study.

| Profile | Strain | Phylogenetic group<sup>a</sup> | Source | References |
| --- | --- | --- | --- | --- |
| Controls | ATCC14579 | IV | Environment | Laboratory collection |
|  | Bt407- | IV | Isolated as a lepidopteran-active strain | [31] |
| | Bt407ΔplcR | IV | Mutant of the Bt407- | [15] |
| Environmental | INRA BC' | II | Vegetable | [20] |
|  | INRA BN | II | Vegetable | [20] |
|  | INRA PF | III | Milk protein | [20] |
|  | INRA A3 | IV | Starch | [20] |
|  | I23 | IV | Cooked apple | [20] |
|  | ATCC10876 | IV | Contaminated flask | Laboratory collection |
|  | SB' | V | Vegetable field | [20] |
|  | I11 | V | Cooked food | [20] |
|  | INRA 1 | VI | Pasteurized zucchini puree | [20] |
|  | INRA BK | VI | Vegetable | [20] |
| Commercial | HD1 | IV | Biopesticide (Dipel®) |  |
|  | ABTS-1857 | IV | Biopesticide (Xentari®) |  |
|  | NB-176 | IV | Biopesticide (Novodor®) |  |
|  | AM65-52 | IV | Biopesticide (Vectobact®) |  |
|  | BCT-7112 | V | Probiotic (Toyocerin®) |  |
| Food Poisoning | NVH0861/00 | II | Ice cream | [20] |
|  | RIVM Bc120 | II | Human feces stools | [32] |
|  | NVH0500/00 | III | Potatoes in cream sauce | [20] |
|  | NVH0075/95 | III | Stew with vegetables | [20] |
|  | NVH1519/00 | III | Stew with deer meat | [20] |
|  | F4433/73 | III | Meat loaf | [20] |
|  | NVH0597/99 | III | Spice mix | [33] |
|  | NVH0230/00 | IV | Mushroom stew | [20] |
|  | NVH1230/88 | IV | Oriental stew | [20] |
|  | F4430/73 | IV | Pea soup | [20] |

<sup>a</sup> Phylogenetic group as described by Guinebretière et al. in 2008 [5].

### 2. Recombinant NheA and SMase proteins and antibody production

#### 2.1. Strain construction

The *B. cereus* group strain NVH0075/95 [24], nowadays classified as *Bacillus paranthracis* (NCBI RefSeq assembly: GCF_027945115.1), was used as DNA source for amplification of *nheA* and *sph* genes. This strain is the reference for Nhe production [22], presents high SMase activity [34], and produces high levels of NheB [9]. Moreover, NheA and SMase protein sequences are highly conserved within the *B. cereus* group strains [4,35]. The sequence coding for the signal peptide, as determined with SignalP (https://services.healthtech.dtu.dk/services/SignalP-6.0/) [36], was removed from the final recombinant sequence. The genomic DNA from the NVH0075/95 strain was obtained using the Puregene Yeast/Bact. kit (Qiagen). Gene sequences were amplified using oligonucleotides 5’-CG**GGATCC**AAGGAGGGGCAAACAGAAGTG-3’ and 5’-AA**CTGCAG**CGTTACTTATTTTTAATGTACTTCAACG-3’ for *nheA* and 5’-CG**GGATCC**GATACGTCTACAGATCAAAACAACATTTTG-3’ and 5’-AA**CTGCAG**GGACTACTTCATAGAAATAGTTGCTGCTAC-3’ for *sph* with Phusion High-Fidelity DNA Polymerase (Thermo Scientific). The amplicons were purified with QIAquick PCR purification kit (Qiagen), then digested with *Bam*HI and *Pst*I restriction enzymes (New England Biolabs), and purified with QIAquick gel extraction kit (Qiagen). Fragments were inserted between *Bam*HI and *Pst*I sites of the plasmid pQE-30 using T4 DNA ligase (New England Biolabs). The resulting sequences enabled the production of proteins with an N-terminal 6 His-Tag. *Escherichia coli* K-12 strain TG1 was transformed by thermal shock with the resultant plasmid constructs separately. Plasmid DNA was prepared using the QIAprep Spin Miniprep Kit (Qiagen) and verified by sequencing (Eurofins). Then, the *E. coli* strain M15 [pREP4] was transformed by thermal shock with verified constructs for recombinant protein production.

#### 2.2. Purification of recombinant proteins

To purify recombinant NheA and SMase proteins, the respective *E. coli* M15 [pREP4] pQE30Ω*nheA* or pQE30Ω*sph* strains were cultivated in LB medium at 30°C, and protein expression was induced with 1mM of IPTG at OD_600_ 0.6. Samples were collected two and four hours after induction for NheA and SMase, respectively, centrifuged at 6000 rpm for 15 min, and stored at-20°C until the following steps. Bacteria were lysed by sonication after a 30 minutes inoculation with 1 mg/mL of lysozyme, centrifuged at 7500 rpm for 20 min at 4 °C. Recombinant proteins were purified from lysate on Ni-NTA resin for 6xHis-tagged protein by following the manufacturer’s recommendations (Qiagen). Purified NheA and SMase were loaded on a 12% SDS-polyacrylamide gel electrophoresis (SDS-PAGE) to verify the size of proteins produced and their purity.

#### 2.3. Antibody production and specificity validation

Rabbit polyclonal antibodies were raised against recombinant NheA and SMase proteins by Covalab (https://www.covalab.com/eu/). The antibodies were purified with Protein A Sepharose 4 Fast Flow (GE Healthcare) and then tested for their specificity using the purified recombinant proteins and the supernatants of NVH0075/95 and ATCC14579 strains by immunoblotting, as described by Candela et al. [37].

### 3. Development of ELISA assay

To detect and quantify NheA and SMase in the supernatant of *B. cereus* group strains, we developed a sandwich Enzyme-Linked Immunosorbent Assay (ELISA) in 96-wells microtiter plates (flat-bottom and high adsorption; Sarstedt). For antigen capture, plates were coated overnight at room temperature with a 100 µL/well solution containing different concentrations of purified anti-SMase or anti-NheA antibodies (capture antibodies), diluted in EPBS (10 mM Sodium phosphate buffer; 150 mM NaCl; pH = 7.2). Then, coated plates were washed three times with ELISA wash buffer (EPBS; 0.1% BSA; 0.05% Tween 20), followed by treatment with ELISA blocking buffer (EPBS; 3% BSA; 0.05% Tween 20) for 1 hour at room temperature with gentle shaking to prevent unspecific binding. After another three-times washing step, plates were set aside to dry at room temperature and stored at 4°C until use. Before use, plates were brought to room temperature. ELISA Dilution buffer (EPBS; 1% BSA; 0.05% Tween 20) was used as a negative control, and serial dilution of purified recombinant proteins was used as a positive control and to construct standard dose-response curves. Controls were added to every single plate. 100 µL/well of each control, and the supernatants of strains Bt407 or Bt407 Δ*plcR* were added to plates and incubated for one hour at room temperature, followed by a three-step washing with ELISA wash buffer. For the detection step, we conjugated HRP to antibodies using the EZ-Link™ Plus Activated Peroxidase (Thermo Scientific) and we tested different HRP-antibody dilutions and time of incubation to determine the optimal assay conditions. The plate was washed again, and 100 µL/well of 1-Step Ultra TMB-ELISA chromogenic substrate (Thermo Scientific) was added. The enzymatic reaction was stopped with 2 M of H_2_SO_4_ (50 µl/well), and the optical density (OD) was measured at 450 nm on a Tecan Infinite® 200 PRO plate reader.

### 4. Quantification of NheA and SMase in the supernatant of *B. cereus* group strains

The amount of NheA and SMase produced by the selected 25 strains of the *B. cereus* group was evaluated from at least three independent cultures of each strain, and samples were analyzed separately for NheA and SMase. Supernatants were diluted in ELISA dilution buffer at ratios ranging from five to 500 times for NheA and from 10 to 200 times for SMase. Diluted supernatants were added to the coated plates in a final volume of 100 µL/well. The standard curves obtained by the serial dilution of the purified recombinant proteins enabled the quantification of both virulence factors in the bacterial supernatants. Each sample was analyzed with at least three technical replicates for each protein.

### 5. Statistical analysis

ANOVA followed by paired t-test was used to compare the production of NheA or SMase among the different profiles or genetic groups (*P* < 0.05). Linear correlation of the average of NheA and SMase production of each strain was analyzed by JMP 9.0 (JMP Statistical discovery LLC). Logistic regression analysis, predictive modeling, threshold determination (Youden index), and the likelihood ratio test were conducted using R-4.5.2 [38], RStudio version 2026.01.1+403 [39], and the packages *pROC* [40] and *glm2* [41]. For these analyses, we considered a binary outcome for the pathogenic profile: nonpathogenic (Environmental and Commercial strain) and pathogenic (Food Poisoning). The predictive modeling considered the combination of both protein titers to predict the outcome and was computed as the Area Under the Curve (AUC), regarding the Receiver operating characteristic (ROC) curve. AUC values can range from 0.5 to 1.0; higher values indicate a better ability of the model to separate the two different profiles. Graphs were plotted with GraphPad Prism version 8.0.2.

## Results

### 1. Production of NheA and SMase recombinant proteins and specificity of anti-NheA and anti-SMase antibodies

Purified recombinant NheA and SMase proteins displayed the expected molecular weights (without the signal peptide and with the 6-His tag) of 42.4 kDa for NheA and 35.6 kDa for SMase, as verified by SDS-PAGE (S1 Fig). These proteins were used for the production of anti-NheA and anti-SMase antibodies. Western blot analysis on supernatant of *B. cereus* group strains NVH0075-95 and ATCC14579 demonstrated the specificity of the purified antibodies against NheA and SMase, as shown in Fig 1A and 1B, respectively. The polyclonal antibodies, anti-NheA and anti-SMase, were selective for the target proteins with the expected molecular weight.

**Fig 1.**
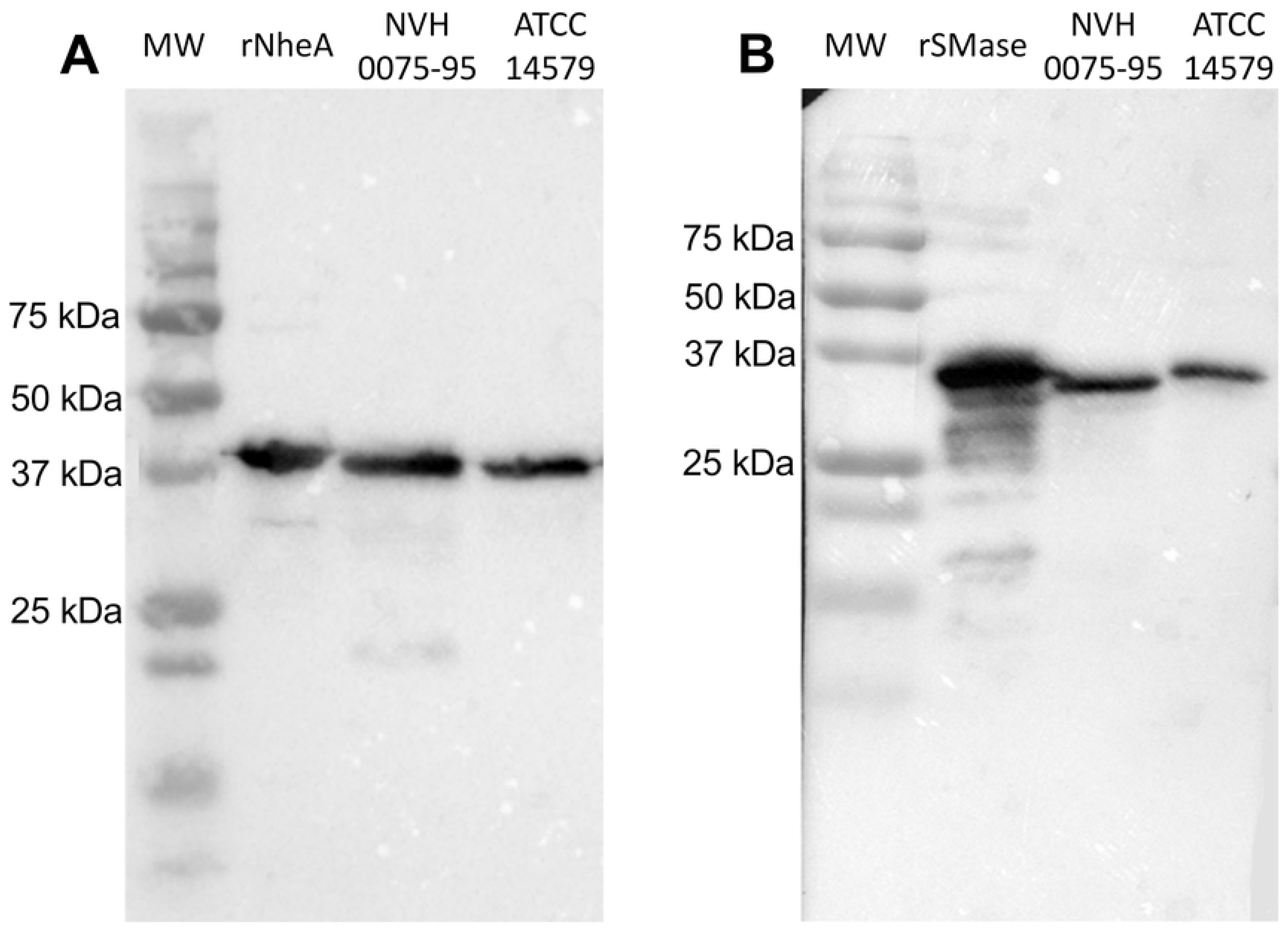
Specificity of Anti-NheA and Anti-SMase antibodies. Western Blot on purified recombinant proteins and on the supernatant of *B. cereus* group strains NVH0075-95 and ATCC14579 cultures obtained two hours after the onset of the stationary phase using polyclonal anti-NheA (A) and anti-SMase (B) antibodies. MW: molecular weight marker.

### 2. Development of an ELISA assay for NheA and SMase immunodetection

The optimal parameters for the development of the ELISA assays were achieved by coating plates with 5 μg/mL of capture antibodies, by using HRP-labeled detection antibodies at 9.13 μg/mL for NheA and 6.23 μg/mL for SMase, with a 45- or 30-min incubation with anti-NheA-HRP or anti-SMase-HRP, respectively, for the detection step. Bt407 Δ*plcR*, which does not produce NheA [13,15], was used as a control strain and showed values (0.174 ± 0.012) comparable to those of the negative control (ELISA dilution buffer; 0.173 ± 0.004). Similar results were observed for SMase detection. Indeed, the supernatant of strain Bt407, which does not produce SMase due to a prophage insertion [42,43], gave similar results as the negative control (0.180 ± 0.006 versus 0.192 ± 0.011, respectively). Taken together, these results demonstrate the specificity of the ELISA assays, with no detectable cross-reactivity against supernatants from strains lacking NheA or SMase production. Then, serial dilutions of purified NheA and SMase were used to obtain the standard dose-response curves shown in Fig 2. These results demonstrate the good sensitivity of our method, showing that the proteins can be detected at concentrations as low as 1 ng/mL for NheA and 0.5 ng/mL for SMase.

**Fig 2.**
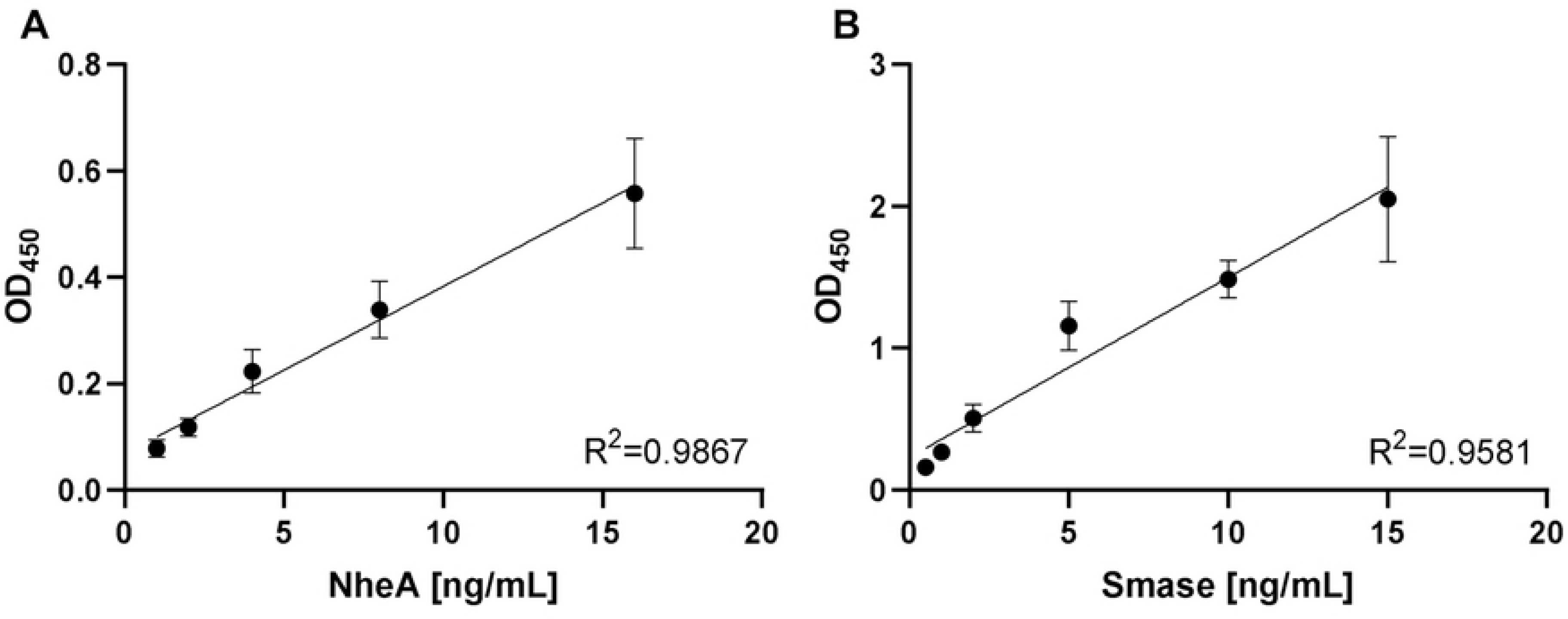
NheA and SMase standard dose-response curves. The OD450 values obtained using the ELISA method were plotted against the doses of NheA and SMase. The regression lines were computed between 1 ng/ml and 16 ng/ml for NheA (A) and between 0.5 ng/ml and 15 ng/ml for SMase (B). Each point is the mean of at least 5 replicates. Error bars represent the standard error of the mean.

### 3. Quantification of NheA production

We quantified NheA in the supernatant of 25 *B. cereus* group strains. NheA titers ranged from 518 ng/mL (strain INRA 1) to 3709 ng/mL (strain NVH0075/95), which represents a more than sevenfold variation among strains (Fig 3A, S1 Table). Strains from the Food Poisoning profile produce approximately twice as much NheA (average of 2244.6 ng/mL) than Environmental (average of 980.8 ng/mL) and Commercial (average of 1155.0 ng/mL) strains (Fig 3B). Moreover, when the NheA quantification was analyzed according to the phylogenetic group of the strains, the results allowed the categorization of these groups into three levels of NheA production (Fig 3C). Strains from group III, almost all Food Poisoning strains, are higher producers of NheA. Strains belonging to groups II and IV, from all the three pathogenic profiles, are intermediate producers. Furthermore, strains from groups V and VI, which are only Environmental and Commercial isolates, are low producers. In summary, the strains that produce the highest levels of NheA are those belonging to both the Food Poisoning profile and to the phylogenetic group III.

**Fig 3.**
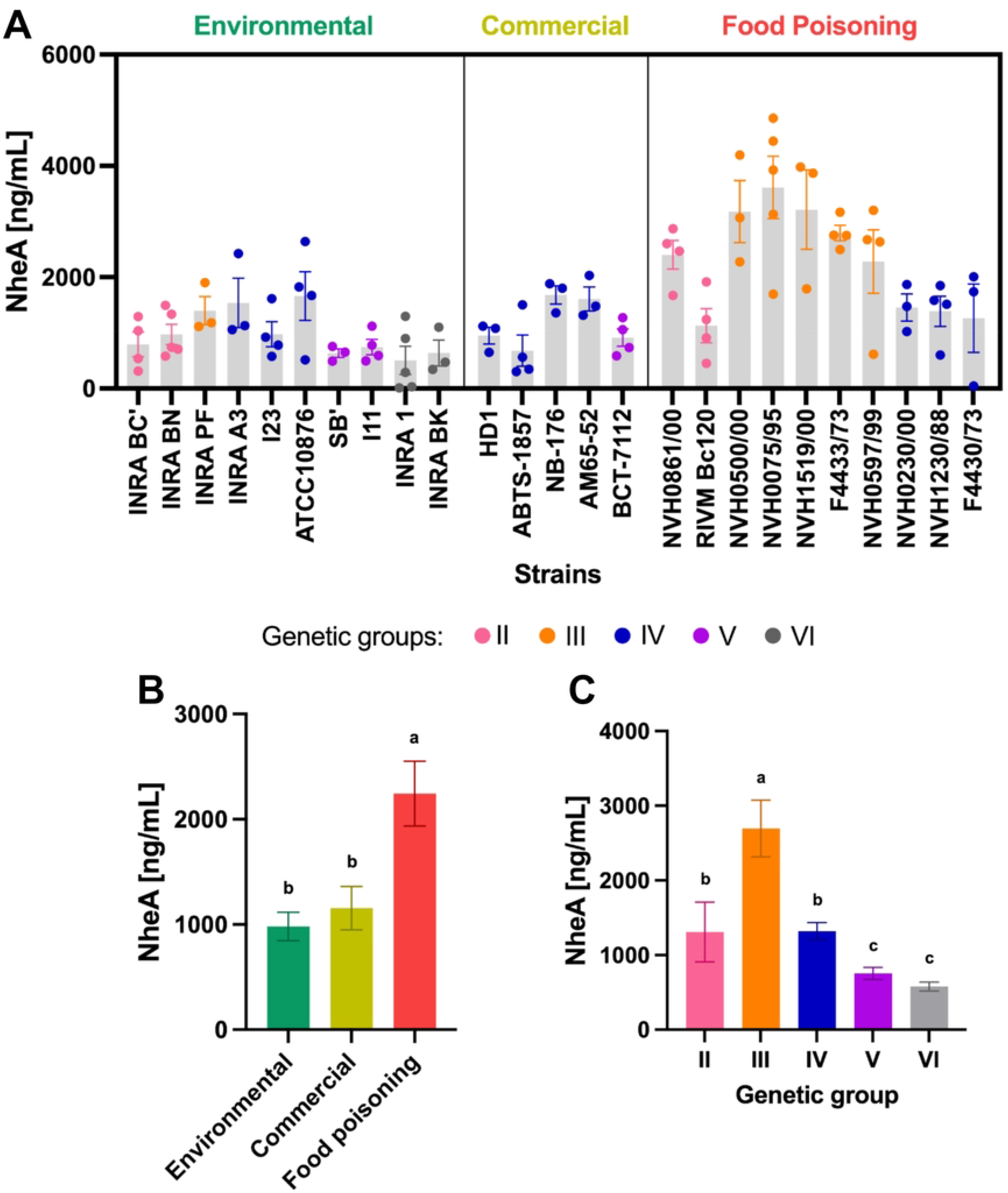
Quantification of NheA production in the supernatant of *B. cereus* group strains. (A) Amount of NheA protein produced by each strain. Grey bars represent the average production of a given strain and each point corresponds to one biological replicate (culture). (B) Average of NheA production by strains from different profiles. (C) Average of NheA production by strains from the different phylogenetic groups. Green: Environmental strains, yellow: Commercial strains, and red: Food Poisoning strains; pink: genetic group II, orange: group III, blue: group IV, purple: group V, grey: group VI. Error bars represent the standard error of the mean. Different letters indicate statistically significant differences in the mean values (P<0.05). Experimental data are detailed in S1 Table.

### 4. Quantification of SMase production

SMase values in the supernatant of the selected strains of the *B. cereus* group (Fig 4A, S1 Table) ranged from non-detectable levels (strain HD1) to 1302.0 ng/mL (strain NVH1519/00). The variation among strains was even greater than for NheA production, with a difference of more than 450-fold when considering the lower detectable titer (2.8 ng/mL for strain AM65-52). On average, Food Poisoning strains produced almost 10 times more SMase than Commercial strains and more than twice as much as Environmental strains, with averages of 707.8 ng/mL, 72.0 ng/mL, and 309.8 ng/mL, respectively (Fig 4B). However, a strain classified as Environmental, isolated from food (milk) and not related to foodborne illness (strain INRA PF), appeared as the second-highest producer of SMase. This strain is classified as belonging to phylogenetic group III that showed a strikingly higher average level of SMase production (947.3 ng/mL) compared to all other groups analyzed in this study: 394.9 ng/mL, 306.7 ng/mL, 226.2 ng/mL, and 112.3 ng/mL, for groups II, IV, V, and VI, respectively (Fig 4C). Similar to the results obtained for NheA, strains from the Food Poisoning profile and phylogenetic group III produce significantly higher levels of SMase than those belonging to other profiles or groups.

**Fig 4.**
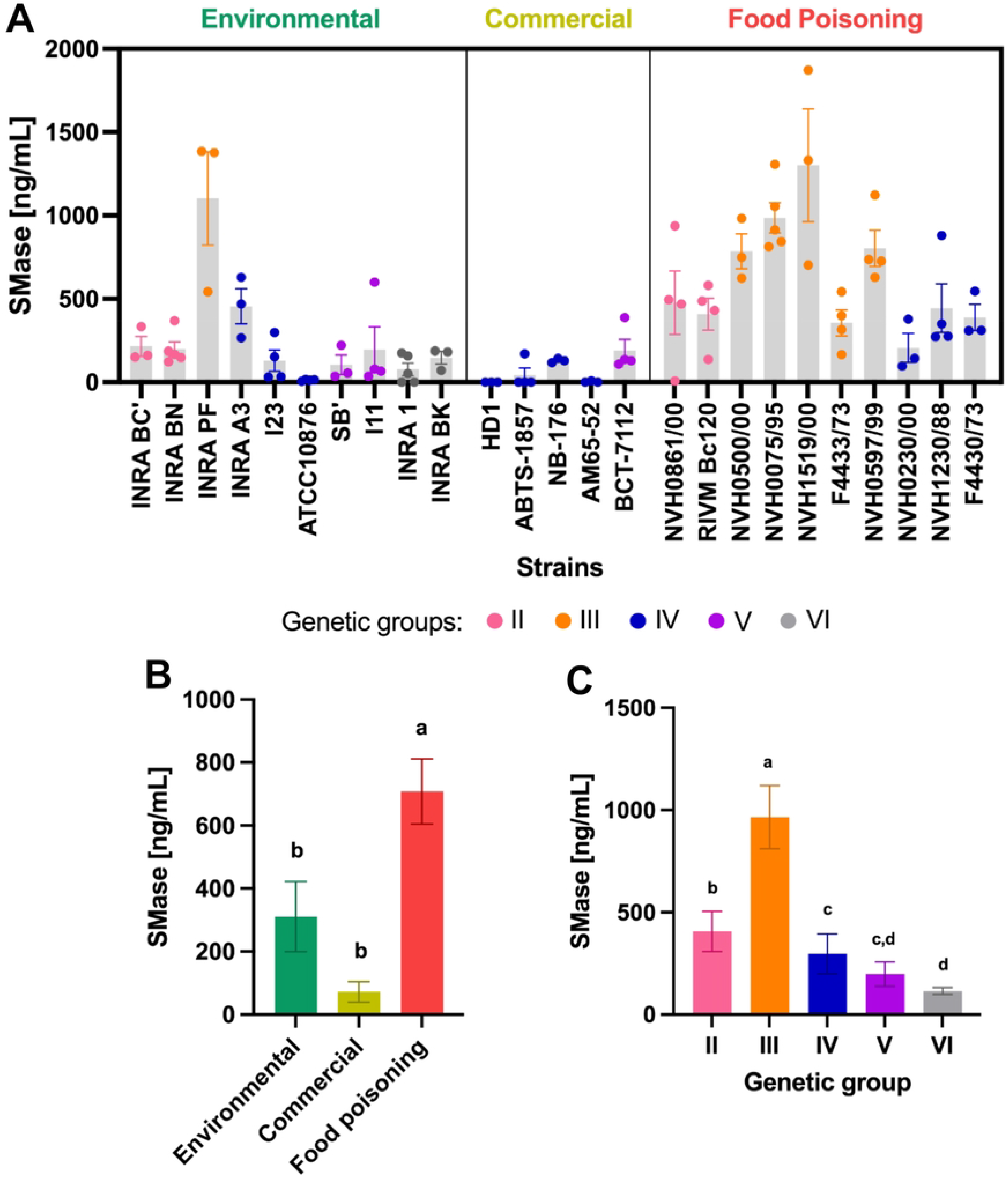
Quantification of SMase production in the supernatant of *B. cereus* group strains. (A) Amount of SMase protein produced by each strain. Grey bars represent the average production of a given strain and each point corresponds to one biological replicate (culture). (B) Average of SMase production by strains from different profiles. (C) Average of SMase production by strains from the different phylogenetic groups. Green: Environmental strains, yellow: Commercial strains, and red: Food Poisoning strains; pink: genetic group II, orange: group III, blue: group IV, purple: group V, grey: group VI. Error bars represent the standard error of the mean. Different letters indicate statistically significant differences in the mean values (P<0.05). Experimental data are detailed in S1 Table.

### 5. Correlation and predictive performance of NheA and SMase production

We performed a combinatorial analysis of NheA and SMase quantification. Firstly, we identified a significant, albeit moderate, positive correlation (*P*<0.01; r^2^=0.45) between the production of these two proteins. Thereafter, we performed a logistic regression analysis to evaluate the correlation between NheA and SMase production and pathogenic profiles. We computed the Receiver operating characteristic (ROC) curve (S2 Fig), showing that the AUC of the NheA model was 0.867, while both the SMase and the combined NheA+SMase models achieved an AUC of 0.907 (Table 2). This analysis indicates that all the three models provide a robust prediction performance to distinguish *B. cereus* diarrheal strains. Furthermore, the likelihood ratio test demonstrated that adding NheA to the SMase model significantly improved the overall fit of the combined model (χ² = 4.97, p = 0.026), suggesting that NheA provides complementary predictive information.

**Table 2.** Predictive performance of NheA and SMase production for discriminating bacterial profiles.

| Model | AUC<br>(95% confidence<br>interval) | Cut-off<br>(ng/mL) | Sensitivity<br>(%) | Specificity<br>(%) | Accuracy<br>(%) |
| --- | --- | --- | --- | --- | --- |
| NheA | 0.867<br>(0.7253 – 1) | 1034.0 | 100 | 66.0 | 80 |
| SMase | 0.907<br>(0.7772 – 1) | 276.5 | 100 | 80.0 | 88 |
| NheA + SMase | 0.907<br>(0.7946 – 1) | - | 100 | 86.6 | 92 |

Then, the protein production thresholds that would best distinguish pathogenic from non-pathogenic strains within our panel were defined from the ROC output using the Youden index. Cutoffs of 1034 ng/mL and 276.5 ng/mL were determined for NheA and SMase, respectively. At these cutoffs, both markers yielded a sensitivity of 100%, while SMase outperformed NheA in terms of specificity (80% versus 66%) and accuracy (80% versus 88%). The combination of the individual NheA and SMase cutoffs achieved a sensitivity of 100%, a specificity of 86.6% and an accuracy of 92%, indicating an improvement in classification performance in comparison with SMase alone (Table 2), as shown with the likelihood ratio. Then, the values of NheA and SMase production for each strain and both individual cutoffs were indicated in a scatter plot (Fig 5). The combined model enables the distinction between Food Poisoning strains and Environmental and Commercial strains, with few exceptions. Indeed, two Environmental isolates, strains INRA A3 and INRA PF, displayed NheA and SMase production profiles similar to those of the Food Poisoning strains. Even if these strains were not isolated from a foodborne outbreak, our results suggest that they should be classified as potentially at risk. Taken together, our results indicate that the combination of NheA and SMase quantifications present a high ability to predict pathogenic profiles, making them valuable potential markers for detecting diarrheal strains.

**Fig 5.**
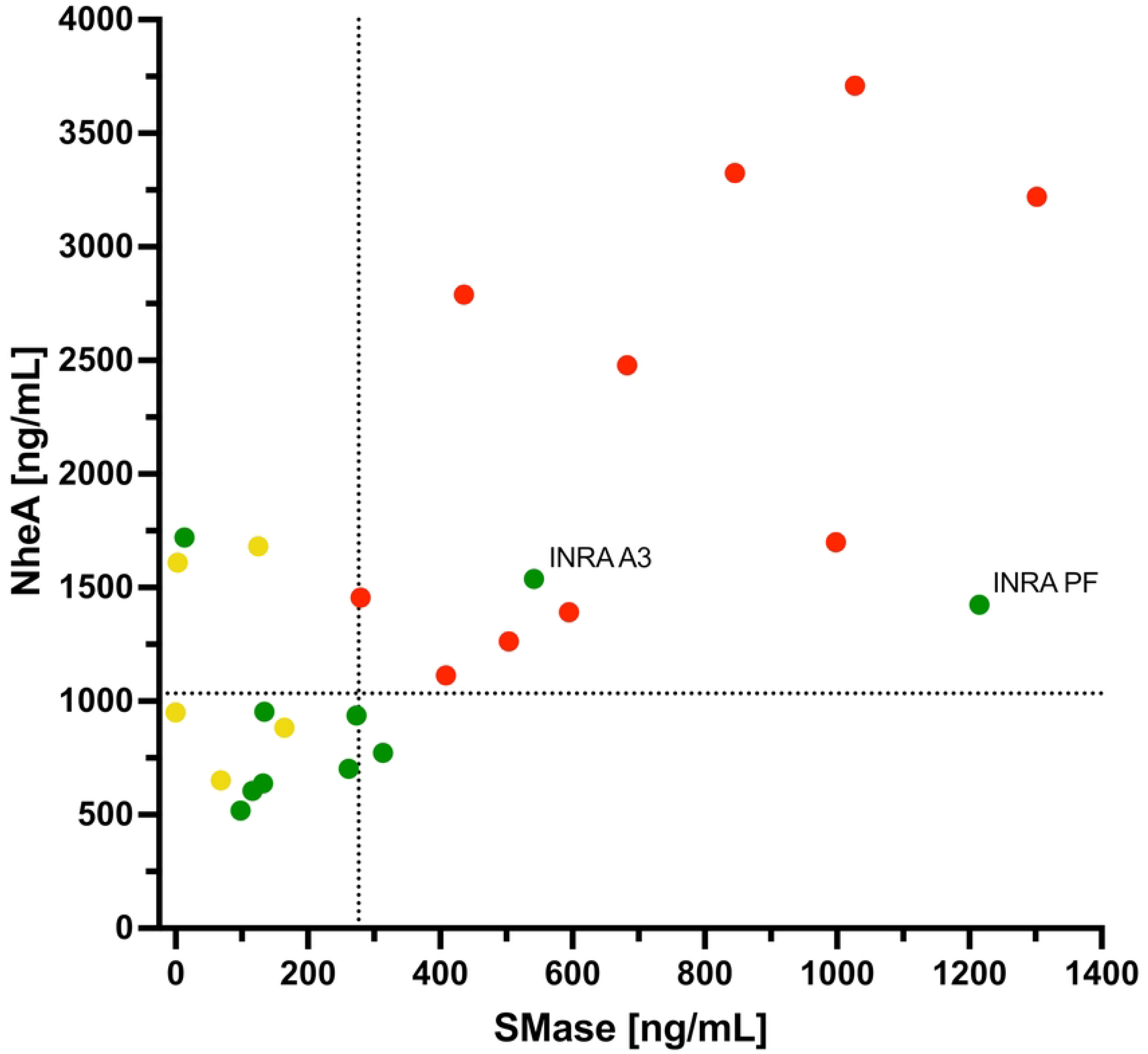
Correlation between NheA and SMase production. Each *B. cereus* group strain is plotted according to NheA and SMase quantification values (in ng/mL). Green: Environmental; yellow: Commercial; and red: Food Poisoning strains. The dotted lines represent the cutoffs: 1034.0 ng/mL for NheA and 276.5 ng/mL for SMase.

## Discussion

Bacteria of the *B. cereus* group are ubiquitous in the environment, including soil, plants, or animals, and can therefore easily contaminate vegetables, milk, or meat. In addition, their spores are resistant to the cleaning and disinfection procedures used in the food industry, such as heat treatments [44]. Furthermore, these bacteria are able to produce dense biofilms, promoting their persistence on food processing equipment in industrial environments [45,46]. Even though most isolates from the *B. cereus* group are innocuous, some are food poisoning agents [2]. Therefore, for economic and health reasons, it is of utmost importance to define a method able to predict the food poisoning potential of unknown food isolates of the *B. cereus* group. In the context of strains producing the emetic toxin or the cytotoxin CytK1, the identification is easily achieved using genetic tools [47,48]. However, the isolates responsible for the diarrheal syndrome do not possess specific genes that are definitively implicated in these cases of food poisoning. Comparative genetic analysis has not been successful in identifying genetic markers to predict the pathogenic potential of these isolates [9]. Indeed, while some genes encoding major virulence factors are present in all strains of the group, others are only sporadically distributed. Consequently, the virulence gene repertoire of a strain does not necessarily correlate with its pathogenic potential [9,17]. Furthermore, the expression of the genes coding for these virulence factors does not fully correlate with their production or pathogenic potential either. Until now, no molecular markers allowed the distinction between potential food poisoning strains and harmless isolates. Another important consideration is that the diarrheic form of *B. cereus* foodborne illness is likely a result of a toxicoinfection [3]. Indeed, the presumed diarrheic virulence factors are susceptible to degradation by food-processing methods and digestive enzymes [49]. The disease is thought to be caused by virulence factors produced in the small intestine of the host by bacteria that were ingested in the form of spores and survived passage through the gastrointestinal tract.

Currently, commercial kits are available to detect *B. cereus* bacteria or their toxins. Some of these are tests that detect the presence of the *B. cereus* bacteria genetic material without assessing its viability, such as the foodproof® SL *Bacillus cereus* Detection Kit (Hygiena). Chromogenic agar media are also available to identify and enumerate viable *B. cereus* bacteria [50], but they do not predict the pathogenic potential of the isolate, as with CHROMagar™ *B. cereus*. Previously, three different kits were available to detect *B. cereus* enterotoxins Nhe and/or Hbl from bacterial culture or directly in food, clinical, or environmental samples [49]. However, these methods detect the toxins either qualitatively (e.g. presence/absence with Duopath) or in a semi-quantitative manner (Via-TECRA and BCET-RPLA Oxoid), which is insufficient to accurately assess the pathogenic potential. Moreover, among these three tests, only BCET-RPLA Oxoid is still available.

Some studies have sought new markers able to reveal the diarrheal pathogenic potential of *B. cereus* group bacteria. Cadot et al. [51] studied virulence factors as candidates for pathogenicity markers. They reported that the *hlyII* gene was not found in the nonpathogenic strains selected for their study, but was present in only 30% of the pathogenic strains, invalidating its use as pathogenicity marker. Jessberger et al. [52] developed a complex scheme to assess the toxic potential of *B. cereus* group isolates, which analyzes spore survival, adhesion and germination, motility, adhesion of vegetative cells, enterotoxin production, and cytotoxicity. Thereafter, they identified a simplified approach for obtaining comparable results. This scheme comprises three different markers: detection of NheB and SMase by slot blot (with monoclonal antibodies), and measurement of exoprotease activity. Kavanaugh et al. [53] described a combination of genetic markers that allow to distinguish clinical (nongastrointestinal infection) isolates, but not food poisoning isolates, from nonpathogenic bacteria. The enteropathogenic potential of *B. cereus* group strains, including Bt-biopesticide strains, was tested *in vitro* and *in vivo* using the Caco-2 cell line and *Drosophila melanogaster* models, respectively [54]. Cytotoxicity and virulence results obtained by these biological models did not correlate with the toxin gene profiles. Finally, the PlcR mutant strain exhibits substantial lower cytotoxic activity on HeLa and CaCo2 cells than the wild-type strain [55]. However, mutant strains independently lacking some PlcR-regulated genes (*hblC*, *cytK-2*, *plcA*, *hlyI*, or *inhA2*) demonstrated no impact in cytotoxicity activity.

In this study, we developed a sandwich ELISA method to quantify NheA and SMase production in the supernatant of *B. cereus* group cultures. We have selected these two PlcR-regulated virulence factors because of their importance in the pathogenesis of the bacteria [21] and the presence of their encoding genes in virtually all *B. cereus* group strains. The amount of secreted NheB enterotoxins was shown to be more reliable to predict pathogenic potential than the genetic traits (presence/absence of the genes or sequence diversity) or the expression of the *nhe* genes [9,56]. However, we decided to use NheA, rather than NheB, as it is the most abundant Nhe component in the supernatant of the type strain ATCC14579 two hours after the onset of the stationary phase [13]. Furthermore, polyclonal anti-NheA and anti-SMase antibodies were used due to their effectiveness at detecting a broader range of protein variants than monoclonal antibodies, such as the slight variations in the studied proteins among strains. Following the optimization process, we have obtained reproducible and sensitive assays with limits of detection of 0.5 ng/mL for SMase and 1 ng/mL for NheA.

The results for NheA and SMase quantification revealed significant variation among individual strains, though we demonstrated that the Food Poisoning strains had significantly higher average production of these two virulence factors than the Environmental or Commercial isolates. We showed a significant positive correlation between NheA and SMase productions, although the strength of the association was moderate. This finding indicates that the two markers share common biological information while retaining substantial independent variability. This moderate correlation may explain why SMase alone exhibited a performance comparable to that of the combined model in terms of AUC, whereas the addition of NheA significantly improved the detection model according to the likelihood ratio test. Together, these results indicate that NheA provides complementary predictive information rather than redundant information, supporting the use of both proteins quantification to improve the classification of bacterial profiles.

The ROC analysis demonstrated that SMase exhibited a higher diagnostic performance than NheA when used as a single marker. The addition of NheA to SMase resulted in a substantial enhancement in the performance of the logistic regression model, while maintaining unaltered AUC. This apparent discrepancy reflects the different aspects assessed by the two statistical approaches. The AUC evaluates the overall discriminative ability of a model, while the likelihood ratio test assesses whether an additional predictor significantly improves model fit. Therefore, although NheA alone showed lower predictive performance than SMase, it provides complementary information that improves discrimination when combined with SMase. These findings indicate that SMase represents the most informative individual marker, while the combined use of both markers improves the reliability of bacterial profile prediction.

Previous studies have shown that the amount of Nhe enterotoxin was higher in food poisoning strains than in those linked to food or the environment [56,57]. However, we highlighted that the difference in SMase production between strains was more outstanding than that observed for NheA. This was particularly evident regarding the commercial strains, which did not produce SMase or produced it at very low levels. This result is in agreement with a previous study showing that biopesticide strains exhibited either no or very low levels of SMase activity [34]. Interestingly, the *sph* gene, which encodes SMase, was found to be disrupted by bacteriophage insertions in some *B. thuringiensis* strains, such as in the HD1 biopesticide strain [42]. This could explain why SMase was not detected in this strain in our assays. The low production of SMase by insecticidal strains may reflect a shift in the pathogenic lifestyle of *B. thuringiensis* bacteria, favoring the insect as their ecological niche [58]. However, due to the close genetic relationship between *B. thuringiensis* and *B. cereus*, and to the high prevalence of enterotoxin genes in commercial *B. thuringiensis* strains it is of growing concern that residues of biopesticides may persist in foodstuffs and, then, be associated with foodborne outbreaks [59,60]. Nevertheless, there is still no direct evidence that Bt-biopesticides lead to diarrheic toxicoinfection [8]. This statement is corroborated by our results regarding low or no SMase production and low or intermediate NheA production in commercial biopesticide strains and by the widespread use of Bt-based products worldwide without a massive causal link to foodborne outbreaks [7].

We demonstrated that strains from phylogenetic group III produced significantly higher concentrations of NheA and SMase than strains from other groups. This finding corroborates the evidence that *B. cereus* group strains belonging to this group could present a higher risk of food poisoning outbreaks [6]. Furthermore, we demonstrated that the combined production levels of these two virulence factors effectively separates Food poisoning strains from Environmental and Commercial isolates. However, the cutoffs values were established based on a representative panel of strains; therefore, they do not constitute absolute thresholds to determine the pathogenic potential of all strains of the *B. cereus* group. Nevertheless, our results point out the applicability of our method, and reveal that even though the INRA A3 and INRA PF strains were isolated from the environment, they could pose a high risk of food poisoning. Statistical analysis confirmed the predictive performance of our method in reliably discriminating between pathogenic and nonpathogenic bacteria within the *B. cereus* group. Therefore, the combined quantification of NheA and SMase is a suitable marker for identifying strains associated with diarrheic food poisoning.

## Conclusion

The accurate identification of *B. cereus* group strains in food products and food-processing environments necessitates reliable tools to assess their potential health risks. This facilitates risk management and reduces the incidence of foodborne outbreaks and associated economic losses. In this study, we developed sandwich ELISA methods for the quantitative determination of NheA and SMase. These markers reliably distinguish diarrheal outbreak-associated isolates from environmental and commercial strains. While SMase proved to be the most informative individual marker, the combined assessment of NheA and SMase yielded complementary predictive information and enhanced the overall classification performance. This approach provides a practical and reliable tool for assessing the diarrheal potential of *B. cereus* group isolates. It could complement current methods used for food safety surveillance and risk assessment.

## Author contribution

**Priscilla F. Cardoso:** Formal analysis, Investigation, Supervision, Validation, Visualization, Writing – original draft, Writing – review and editing. **Nathalie Gilois:** Investigation, Methodology, Validation, Visualization, Writing – original draft, Writing – review and editing. **Gislayne T. Vilas-Bôas:** Funding acquisition, Writing – review and editing. **Didier Lereclus:** Conceptualization, Project administration, Validation, Writing – review and editing. **Michel Gohar:** Conceptualization, Formal analysis, Funding acquisition, Methodology, Project administration, Supervision, Validation, Writing – review and editing. **Stéphane Perchat:** Formal analysis, Supervision, Validation, Visualization, Writing – review and editing. **Leyla Slamti:** Supervision, Validation, Writing – review and editing.

## Acknowledgements

This work was supported by the European Union’s Horizon 2020 research and innovation programme under Grant Agreement No 773830 (Toxdetect project). Cardoso, P.F. was supported by a fellowship from National Council for Scientific and Technological Development – CNPq. We thank Rebecca Demonti for technical assistance. We also thank Véronique Broussolle and Ole Andreas Økstad for providing some strains.

## Supporting information

**S1 Fig. Production of recombinant NheA and SMase proteins.** (A) rNheA and (B) rSMase SDS-PAGE gel with 3.6μg of purified recombinant proteins obtained from *E. coli*. Gels were cropped to retain the relevant wells. MW: molecular weight marker.

**S1 Table. NheA and SMase quantification in culture supernatants of *Bacillus cereus* group strains.**

**S2 Fig. Receiver operating characteristic curve (ROC).** ROC curves for prediction of pathogenic profile (food poisoning vs. non-pathogenic) are based on protein levels of NheA, SMase and combined NheA + SMase in the supernatant of *B. cereus* strains measured by ELISA.

## References

1. Bottone EJ. *Bacillus cereus*, a volatile human pathogen. Clin Microbiol Rev. 2010;23: 382–398. doi:10.1128/CMR.00073-09

2. Ehling-Schulz M, Lereclus D, Koehler TM. The *Bacillus cereus* group: *Bacillus* species with pathogenic potential. Microbiol Spectr. 2019;7 (3): GPP3-0032-2018. doi:10.1128/microbiolspec.GPP3-0032-2018

3. Jovanovic J, Ornelis VFM, Madder A, Rajkovic A. *Bacillus cereus* food intoxication and toxicoinfection. Compr Rev Food Sci Food Saf. 2021;20: 3719–3761. doi:10.1111/1541-4337.12785

4. Stenfors Arnesen LP, Fagerlund A, Granum PE. From soil to gut: *Bacillus cereus* and its food poisoning toxins. FEMS Microbiol Rev. 2008;32: 579–606. doi:10.1111/j.1574-6976.2008.00112.x

5. Guinebretière M-H, Thompson FL, Sorokin A, Normand P, Dawyndt P, Ehling-Schulz M, et al. Ecological diversification in the *Bacillus cereus* Group. Environ Microbiol. 2008;10: 851–865. doi:10.1111/j.1462-2920.2007.01495.x

6. Guinebretière MH, Velge P, Couvert O, Carlin F, Debuyser ML, Nguyen-The C. Ability of *Bacillus cereus* group strains to cause food poisoning varies according to phylogenetic affiliation (groups I to VII) rather than species affiliation. J Clin Microbiol. 2010;48: 3388–3391. doi:10.1128/JCM.00921-10

7. Biggel M, Jessberger N, Kovac J, Johler S. Recent paradigm shifts in the perception of the role of *Bacillus thuringiensis* in foodborne disease. Food Microbiol. 2022;105: 104025. doi:10.1016/j.fm.2022.104025

8. Slamti L, Lereclus D. *Bacillus thuringiensis* and insects: a century of intimate history. J Bacteriol. 2026;208: 1–18. doi:10.1128/jb.00381-25

9. Jessberger N, Krey VM, Rademacher C, Böhm ME, Mohr AK, Ehling-Schulz M, et al. From genome to toxicity: A combinatory approach highlights the complexity of enterotoxin production in *Bacillus cereus*. Front Microbiol. 2015;6: 1–15. doi:10.3389/fmicb.2015.00560

10. Agaisse H, Gominet M, Okstad OA, Kolsto A-B, Lereclus D. PlcR is a pleiotropic regulator of extracellular virulence factor gene expression in *Bacillus thuringiensis*. Mol Microbiol. 1999;32: 1043–1053. doi:10.1046/j.1365-2958.1999.01419.x

11. Slamti L, Lereclus D. A cell-cell signaling peptide activates the PlcR virulence regulon in bacteria of the *Bacillus cereus* group. EMBO J. 2002;21: 4550–9. Available: http://www.ncbi.nlm.nih.gov/pubmed/12198157

12. Gohar M, Faegri K, Perchat S, Ravnum S, Økstad OA, Gominet M, et al. The PlcR Virulence Regulon of *Bacillus cereus*. Yang C-H, editor. PLoS One. 2008;3: e2793. doi:10.1371/journal.pone.0002793

13. Gohar M, Økstad OA, Gilois N, Sanchis V, Kolstø AB, Lereclus D. Two-dimensional electrophoresis analysis of the extracellular proteome of *Bacillus cereus* reveals the importance of the PlcR regulon. Proteomics. 2002;2: 784–791. doi:10.1002/1615-9861(200206)2:6<784::AID-PROT784>3.0.CO;2-R

14. Callegan MC, Kane ST, Cochran DC, Gilmore MS, Gominet M, Lereclus D. Relationship of *plcR*-Regulated Factors to *Bacillus* Endophthalmitis Virulence. Infect Immun. 2003;71: 3116–3124. doi:10.1128/IAI.71.6.3116-3124.2003

15. Salamitou S, Ramisse F, Brehelin M, Bourguet D, Gilois N, Gominet M, et al. The *plcR* regulon is involved in the opportunistic properties of *Bacillus thuringiensis* and *Bacillus cereus* in mice and insects. Microbiology (N Y). 2000;146: 2825–2832. doi:10.1099/00221287-146-11-2825

16. Jessberger N, Rademacher C, Krey VM, Dietrich R, Mohr AK, Böhm ME, et al. Simulating intestinal growth conditions enhances toxin production of enteropathogenic Bacillus cereus. Front Microbiol. 2017;8. doi:10.3389/fmicb.2017.00627

17. Dietrich R, Jessberger N, Ehling-Schulz M, Märtlbauer E, Granum PE. The Food Poisoning Toxins of *Bacillus cereus*. Toxins (Basel). 2021;13. doi:10.3390/toxins13020098

18. Fagerlund A, Lindbäck T, Granum PE. *Bacillus cereus* cytotoxins Hbl, Nhe and CytK are secreted via the Sec translocation pathway. BMC Microbiol. 2010;10. doi:10.1186/1471-2180-10-304

19. Prince C, Kovac J. Regulation of Enterotoxins Associated with *Bacillus cereus Sensu Lato* Toxicoinfection. Appl Environ Microbiol. 2022;88: 1–12. doi:10.1128/aem.00405-22

20. Kamar R, Gohar M, Jéhanno I, Réjasse A, Kallassy M, Lereclus D, et al. Pathogenic potential of *Bacillus cereus* strains as revealed by phenotypic analysis. J Clin Microbiol. 2013;51: 320–323. doi:10.1128/JCM.02848-12

21. Doll VM, Ehling-Schulz M, Vogelmann R. Concerted Action of Sphingomyelinase and Non-Hemolytic Enterotoxin in Pathogenic *Bacillus cereus*. PLoS One. 2013;8. doi:10.1371/journal.pone.0061404

22. Granum PE, O’sullivan K, Lund T. The sequence of the non-haemolytic enterotoxin operon from *Bacillus cereus*. FEMS Microbiol Lett. 1999;177: 225–229. doi:10.1111/j.1574-6968.1999.tb13736.x

23. Lindbäck T, Fagerlund A, Rødland MS, Granum PE. Characterization of the *Bacillus cereus* Nhe enterotoxin. Microbiology (N Y). 2004;150: 3959–3967. doi:10.1099/mic.0.27359-0

24. Lund T, Granum PE. Characterisation of a non-haemolytic enterotoxin complex from *Bacillus cereus* isolated after a foodborne outbreak. FEMS Microbiol Lett. 1996;141: 151–156. doi:10.1016/0378-1097(96)00208-X

25. Lindbäck T, Hardy SP, Dietrich R, Sødring M, Didier A, Moravek M, et al. Cytotoxicity of the *Bacillus cereus* Nhe Enterotoxin Requires Specific Binding Order of Its Three Exoprotein Components. Infect Immun. 2010;78: 3813–3821. doi:10.1128/IAI.00247-10

26. Oda M, Hashimoto M, Takahashi M, Ohmae Y, Seike S, Kato R, et al. Role of sphingomyelinase in infectious diseases caused by *Bacillus cereus*. PLoS One. 2012;7: 1–11. doi:10.1371/journal.pone.0038054

27. Gilmore MS, Cruz-Rodz AL, Leimeister-Wachter M, Kreft J, Goebel W. A *Bacillus cereus* cytolytic determinant, cereolysin AB, which comprises the phospholipase C and sphingomyelinase genes: Nucleotide sequence and genetic linkage. J Bacteriol. 1989;171: 744–753. doi:10.1128/jb.171.2.744-753.1989

28. Beecher DJ, Wong ACL. Cooperative, synergistic and antagonistic haemolytic interactions between haemolysin BL, phosphatidylcholine phospholipase C and sphingomyelinase from *Bacillus cereus*. Microbiology (N Y). 2000;146: 3033–3039. doi:10.1099/00221287-146-12-3033

29. Flores-Díaz M, Monturiol-Gross L, Naylor C, Alape-Girón A, Flieger A. Bacterial Sphingomyelinases and Phospholipases as Virulence Factors. Microbiology and Molecular Biology Reviews. 2016;80: 597–628. doi:10.1128/mmbr.00082-15

30. Gilois N, Ramarao N, Bouillaut L, Perchat S, Aymerich S, Nielsen-LeRoux C, et al. Growth-related variations in the *Bacillus cereus* secretome. Proteomics. 2007;7: 1719–1728. doi:10.1002/pmic.200600502

31. Lereclus D, Arantès O, Chaufaux J, Lecadet M. Transformation and expression of a cloned delta-endotoxin gene in *Bacillus thuringiensis*. FEMS Microbiol Lett. 1989;51: 211–7. Available: http://www.ncbi.nlm.nih.gov/pubmed/2550317

32. Carlin F, Albagnac C, Rida A, Guinebretière MH, Couvert O, Nguyen-the C. Variation of cardinal growth parameters and growth limits according to phylogenetic affiliation in the Bacillus cereus Group. Consequences for risk assessment. Food Microbiol. 2013;33: 69–76. doi:10.1016/j.fm.2012.08.014

33. Tourasse NJ, Kolstø AB. Survey of group I and group II introns in 29 sequenced genomes of the Bacillus cereus group: Insights into their spread and evolution. Nucleic Acids Res. 2008;36: 4529–4548. doi:10.1093/nar/gkn372

34. Johler S, Kalbhenn EM, Heini N, Brodmann P, Gautsch S, Bagcioglu M, et al. Enterotoxin production of *Bacillus thuringiensis* isolates from biopesticides, foods, and outbreaks. Front Microbiol. 2018;9: 1–11. doi:10.3389/fmicb.2018.01915

35. Böhm ME, Huptas C, Krey VM, Scherer S. Massive horizontal gene transfer, strictly vertical inheritance and ancient duplications differentially shape the evolution of *Bacillus cereus* enterotoxin operons *hbl*, *cytK* and *nhe*. BMC Evol Biol. 2015;15: 1–17. doi:10.1186/s12862-015-0529-4

36. Almagro Armenteros JJ, Tsirigos KD, Sønderby CK, Petersen TN, Winther O, Brunak S, et al. SignalP 5.0 improves signal peptide predictions using deep neural networks. Nat Biotechnol. 2019;37: 420–423. doi:10.1038/s41587-019-0036-z

37. Candela T, Fagerlund A, Buisson C, Gilois N, Kolstø AB, Økstad OA, et al. CalY is a major virulence factor and a biofilm matrix protein. Mol Microbiol. 2019;111: 1416– 1429. doi:10.1111/mmi.14184

38. R Core Team. R: A language and environment for statistical computing. Vienna, Austria: R Foundation for Statistical Computing; 2021. Available: https://www.r-project.org/

39. Posit team. RStudio: Integrated Development Environment for R. Boston, MA: Posit Software, PBC; 2025. Available: http://www.posit.co/

40. Robin X, Turck N, Hainard A, Tiberti N, Lisacek F, Sanchez J-C, et al. pROC: an open-source package for R and S+ to analyze and compare ROC curves. BMC Bioinformatics. 2011;12: 77. doi:10.1186/1471-2105-12-77

41. Marschner, Ian C. glm2: Fitting Generalized Linear Models with Convergence Problems. R J. 2011;3: 12. doi:10.32614/RJ-2011-012

42. Biggel M, Etter D, Corti S, Brodmann P, Stephan R, Ehling-Schulz M, et al. Whole Genome Sequencing Reveals Biopesticidal Origin of *Bacillus thuringiensis* in Foods. Front Microbiol. 2022;12. doi:10.3389/fmicb.2021.775669

43. Sheppard AE, Poehlein A, Rosenstiel P, Liesegang H, Schulenburg H. Complete Genome Sequence of *Bacillus thuringiensis* Strain 407 Cry-. Genome Announc. 2013;1: 7–8. doi:10.1128/genomeA.00158-12

44. Faille C, Fontaine F, Bénézech T. Potential occurrence of adhering living Bacillus spores in milk product processing lines. J Appl Microbiol. 2001;90: 892–900.

45. Lequette Y, Garénaux E, Tauveron G, Dumez S, Perchat S, Slomianny C, et al. Role played by exosporium glycoproteins in the surface properties of *Bacillus cereus* spores and in their adhesion to stainless steel. Appl Environ Microbiol. 2011;77: 4905–4911. doi:10.1128/AEM.02872-10

46. Majed R, Faille C, Kallassy M, Gohar M. *Bacillus cereus* Biofilms - same, only different. Front Microbiol. 2016;7: 1–16. doi:10.3389/fmicb.2016.01054

47. Ehling-Schulz M, Fricker M, Scherer S. Identification of emetic toxin producing *Bacillus cereus* strains by a novel molecular assay. FEMS Microbiol Lett. 2004;232: 189–195. doi:10.1016/S0378-1097(04)00066-7

48. Guinebretière M-H, Fagerlund A, Granum PE, Nguyen-The C. Rapid discrimination of cytK-1 and cytK-2 genes in Bacillus cereus strains by a novel duplex PCR system. FEMS Microbiol Lett. 2006;259: 74–80. doi:10.1111/j.1574-6968.2006.00247.x

49. Ceuppens S, Rajkovic A, Hamelink S, Van de Wiele T, Boon N, Uyttendaele M. Enterotoxin Production by *Bacillus cereus* Under Gastrointestinal Conditions and Their Immunological Detection by Commercially Available Kits. Foodborne Pathog Dis. 2012;9: 1130–1136. doi:10.1089/fpd.2012.1230

50. Fuchs E, Raab C, Brugger K, Ehling-Schulz M, Wagner M, Stessl B. Performance Testing of *Bacillus cereus* Chromogenic Agar Media for Improved Detection in Milk and Other Food Samples. Foods. 2022;11. doi:10.3390/foods11030288

51. Cadot C, Tran SL, Vignaud ML, De Buyser ML, Kolstø AB, Brisabois A, et al. InhA1, NprA, and HlyII as candidates for markers to differentiate pathogenic from nonpathogenic *Bacillus cereus* strains. J Clin Microbiol. 2010;48: 1358–1365. doi:10.1128/JCM.02123-09

52. Jessberger N, Kranzler M, Da Riol C, Schwenk V, Buchacher T, Dietrich R, et al. Assessing the toxic potential of enteropathogenic *Bacillus cereus*. Food Microbiol. 2019;84: 103276. doi:10.1016/j.fm.2019.103276

53. Kavanaugh DW, Glasset B, Dervyn R, Guérin C, Plancade S, Herbin S, et al. New genetic biomarkers to differentiate non-pathogenic from clinically relevant *Bacillus cereus* strains. Clinical Microbiology and Infection. 2022;28: 137.e1–137.e8. doi:10.1016/j.cmi.2021.05.035

54. Fichant A, Lanceleur R, Hachfi S, Brun-Barale A, Blier AL, Firmesse O, et al. New Approach Methods to Assess the Enteropathogenic Potential of Strains of the *Bacillus cereus* Group, including *Bacillus thuringiensis*. Foods. 2024;13: 1–19. doi:10.3390/foods13081140

55. Ramarao N, Lereclus D. Adhesion and cytotoxicity of *Bacillus cereus* and *Bacillus thuringiensis* to epithelial cells are FlhA and PlcR dependent, respectively. Microbes Infect. 2006;8: 1483–1491. doi:10.1016/j.micinf.2006.01.005

56. Moravek M, Dietrich R, Buerk C, Broussolle V, Guinebretière MH, Granum PE, et al. Determination of the toxic potential of *Bacillus cereus* isolates by quantitative enterotoxin analyses. FEMS Microbiol Lett. 2006;257: 293–298. doi:10.1111/j.1574-6968.2006.00185.x

57. Guinebretière MH, Broussolle V, Nguyen-The C. Enterotoxigenic profiles of food-poisoning and food-borne *Bacillus cereus* strains. J Clin Microbiol. 2002;40: 3053– 3056. doi:10.1128/JCM.40.8.3053-3056.2002

58. Jensen GB, Hansen BM, Eilenberg J, Mahillon J. The hidden lifestyles of *Bacillus cereus* and relatives. Environ Microbiol. 2003;5: 631–40. doi:10.1046/j.1462-2920.2003.00461.x

59. Bonis M, Felten A, Pairaud S, Dijoux A, Maladen V, Mallet L, et al. Comparative phenotypic, genotypic and genomic analyses of *Bacillus thuringiensis* associated with foodborne outbreaks in France. PLoS One. 2021;16: 1–23. doi:10.1371/journal.pone.0246885

60. EFSA. Risks for public health related to the presence of *Bacillus cereus* and other *Bacillus* spp. including *Bacillus thuringiensis* in foodstuffs. EFSA Journal. 2016;14. doi:10.2903/j.efsa.2016.4524

